# Computational model of 3D cell migration based on the molecular clutch mechanism

**DOI:** 10.1101/2021.09.29.462287

**Authors:** Samuel Campbell, Rebecca Zitnay, Michelle Mendoza, Tamara C Bidone

## Abstract

The external environment is a regulator of cell activity. Its stiffness and microstructure can either facilitate or prevent 3D cell migration in both physiology and disease. 3D cell migration results from force feedbacks between the cell and the extracellular matrix (ECM). Adhesions regulate these force feedbacks by working as molecular clutches that dynamically bind and unbind the ECM. Because of the interdependency between ECM properties, adhesion dynamics, and cell contractility, how exactly 3D cell migration occurs in different environments is not fully understood. In order to elucidate the effect of ECM on 3D cell migration through force-sensitive molecular clutches, we developed a computational model based on a lattice point approach. Results from the model show that increases in ECM pore size reduce cell migration speed. In contrast, matrix porosity increases it, given a sufficient number of ligands for cell adhesions and limited crowding of the matrix from cell replication. Importantly, these effects are maintained across a range of ECM stiffnesses’, demonstrating that mechanical factors are not responsible for how matrix microstructure regulates cell motility.

## Introduction

The ability of cells to adapt to and migrate through 3D matrices is essential for physiological and pathological processes, including morphogenesis during development, wound healing, tissue generation, and cancer invasion. Naturally adherent cells, such as tissue fibroblasts, epithelial cells, and cancer cells, sense and respond to their matrix through adhesions (1). Adhesions transmit force bidirectionally between the contractile actomyosin cytoskeleton and the external matrix. Understanding the interplay between matrix properties, cell adhesion, and motility is important for our understanding of cell behavior in physiology and disease, for engineering scaffolds for tissue regeneration, and the future development of therapies for cancer cell invasion.

The microarchitecture of the extracellular matrix (ECM), including pore size and porosity, can either promote or inhibit 3D cell motility. Dense and loose ECM networks can impose steric hindrances on cell migration (1). In collagen hydrogels with pores of 1-20 μm, reducing the size of matrix pores limits cell migration speed of human fibrosarcoma and breast cancer cells, regardless of stiffness (2–4). In contrast, pore size is inversely correlated with cell migration for larger pores sizes. Mouse fibroblasts migrate faster on 3D collagen-glycosaminoglycan scaffolds with pores of 100 μm than in scaffolds with larger pores (5). Similarly, human fibroblasts move faster in more porous 3D silk fibroin scaffolds with smaller pores of 100 μm than in less porous scaffolds with larger pores of 150 μm or 250 μm (6). In the former case, the smallest pore sizes limit 3D migration because they constrain nuclear passage (3). In these latter cases, smaller well-connected pores in porous matrices can promote 3D cell migration by providing sufficient area for adhesion formation, as also suggested in other studies (5, 7).

3D cell migration requires adhesions to the ECM (8, 9). Adhesion molecules, i.e., integrins, sense and respond to forces by changing ligand-bound stability. Integrins adhesions behave as force-sensitive molecular clutches, ensuring bidirectional force transmission between cells and ECM when the clutches are engaged and no force transmission when they are disengaged (10–14). This engagement and disengagement of the clutches depend on the force on them, which in turn depends on the difference between actomyosin contractility and external stiffness.

The mechanical properties of the ECM also affect 3D cell migration. Cells tune their contractility to the surrounding ECM and migrate toward stiffer substrate regions in a process termed durotaxis (10, 15–18). Fibroblasts, mesenchymal stem cells, and cancer cells migrating in collagen on top of polyacrylamide gels of 1-65 kPa exhibit increased migration velocity on stiffer substrates. Similarly, mesenchymal stem cells migrating in 3D gels of 1-40 kPa migrate towards the stiffer gel area (17). A biphasic relationship between migration speed and matrix stiffness was also observed for breast carcinoma cells and human glioma cells moving in 3D (19–21). In addition, fibroblasts and endothelial cells move faster in stiff 3D collagen gels with greater porosity than in soft ECM with less porosity (22). However, resolving the effects of ECM microarchitecture and stiffness has been challenged by the limited ability to *in vitro* manipulate one parameter without the others.

Because of the difficulty in experimentally isolating the effects of ECM microstructure and mechanics on cell migration and monitoring the dynamics of cell adhesions in 3D, how exactly 3D environments regulate cell migratory behavior remains largely unknown. This study aims to elucidate how cells move in 3D matrices through contractility adaptation of integrin-based molecular clutches. For this, we developed a novel 3D model based on a lattice point approach. In the model, matrix pores are explicit 3D regions of defined volumes, and cells are spherical particles that create dynamic integrin-based adhesions. Cell displacement within the model is dependent on three components: 1) the force feedback between the cell and the ECM, 2) the contact surface area between cells and ECM pores, and 3) the availability of ligands, which regulates adhesions dynamics; over time, these cell displacements are monitored and the relations between properties of the ECM and cell velocity analyzed. Our results show a biphasic relation between cell motility and matrix stiffness, consistent with a range of previous studies of mechanosensitive cell motility (23–25). Furthermore, this relation exists in environments with different microarchitectures, supporting the general idea that the pore size and porosity of a 3D ECM regulate variations in cell motion, independent of mechanical factors.

## Methods

In order to isolate the contributions of environment microstructure from mechanics and study their effects on 3D cell migration, we developed and implemented a computational model based on a lattice point approach (26–28). Unlike previous 3D cell migration models and lattice point simulation methods, we incorporated the dynamics of adhesions clutches that respond to force feedback between cell contractility and external matrix properties. A molecular-clutch based mechanism for 3D cell migration was previously reported for fibroblasts’ in 3D collagen and fibrin matrices (22, 29, 30). Our model evaluates the available area for cell adhesions, ligands concentration, the difference between actomoyosin contractility and matrix stiffness, and their effects on the stability of molecular clutches to govern cell motion. The stability of molecular clutches determines adhesions number, which, in turn, regulates cell displacement, accounting for local crowding. The overall 3D simulation domain is a cube with sides of 1.5 mm, divided into a regular grid (lattice), with each lattice point representing a pore for cells to occupy. Each pore is spherical and provides the space and surface where cells move to and create adhesions for force transmission and cell speed regulation (Fig. 1A). Individual cells are spherical, with a defined volume and corresponding surface area.

**Figure 1.**
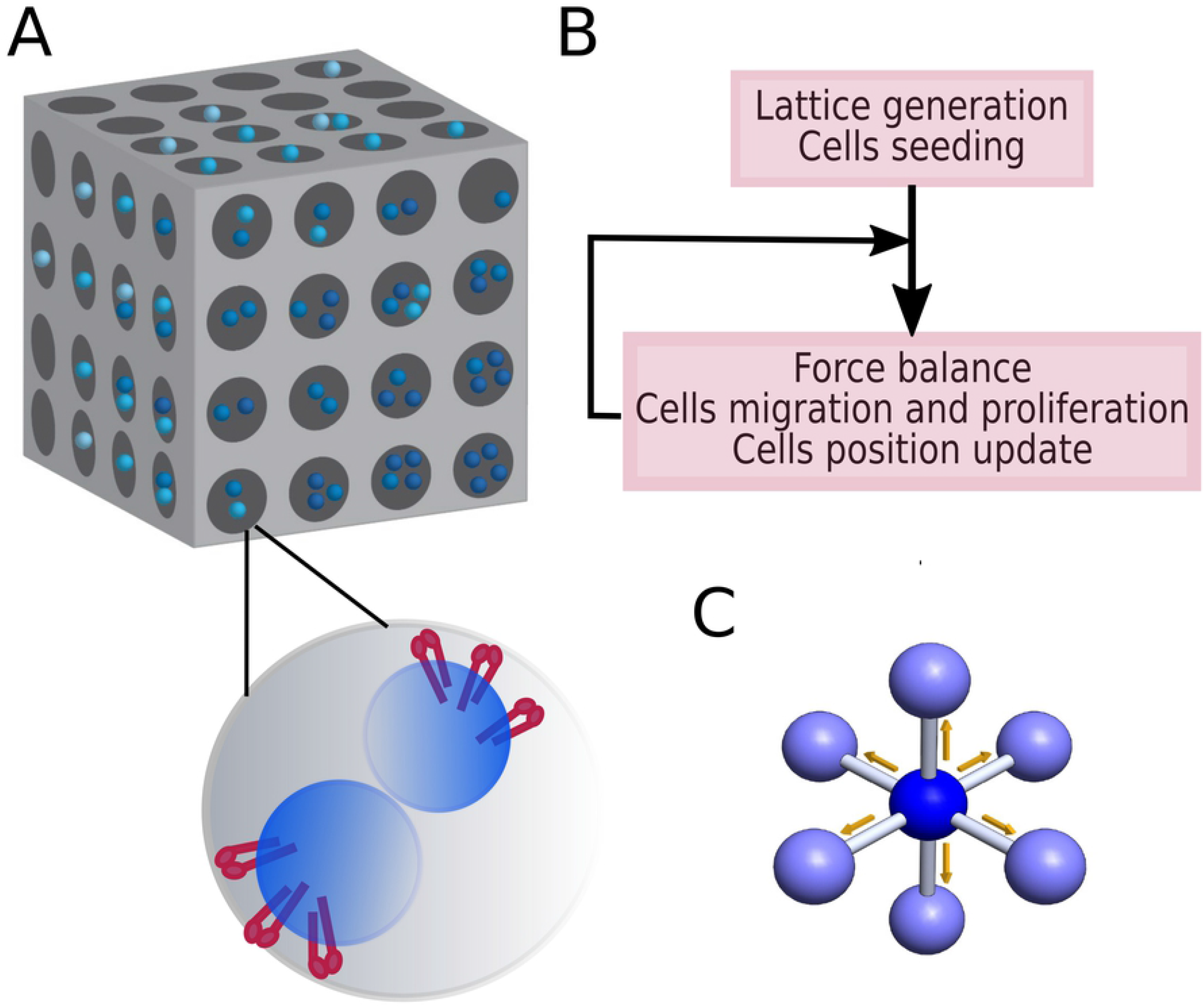
Lattice point model of cell migration in a 3D matrix. **A**. Schematic representation of the computational domain, with lattice points mimicking matrix pores of homogeneous dimensions. Zoomed section represents one lattice point, with two spherical cells (in blue) adhering to the pore surface through integrin adhesions (in red). **B**. Flowchart of the model. After initializing the computational domain, the model parameters, and the cell density, the algorithm iteratively updates cell positions as a function of force balance between internal contractility, matrix stiffness, number of ligated integrins, *n*_*bonds*_, and external viscosity. **C**. Each cell randomly moves in one of the six directions, in one of the neighboring lattice points, provided that it is not filled.

### Simulation domain and main algorithmic steps

In the model, cells move throughout the 3D matrix and replicate. First, a random direction for the cell’s move is chosen. Then, the model predicts the cell migration distance assuming a net force balance between the cell traction, *F*_trac_, the formation and disassembly of integrin adhesions with matrix ligands, and the environment drag, *F*_drag_, due to the viscous resistance of the 3D matrix. Last, based on cell migration distance and local crowding, cell displacement can occur. At each time step of the simulations, the program keeps track of cell positions (Fig. 1B), free volume in each pore, and the contact area between each cell and its pore to determine adhesion number and total cell count.

Cells can move randomly in one of six directions (Fig. 1C), each corresponding to one neighboring lattice point (neighboring pore). If the pore is full, a new random direction is chosen. In runs in which cell proliferation occurs, new cells are also generated, following migration. Proliferation depends on the input proliferation rate which determines the proliferation probability (Fig. 1B). In this case, cells are classified by generation number, with “1” indicating seeded cells, “2” indicating cells generated from generation 1 and so on.

### Force feedbacks between cells and the 3D matrix determine cells migration

At every timestep of the simulation, the net force acting on each cell is calculated to determine cell migration as a function of time. The net force acting on each cell depends on cell traction, number of adhesions, and environment drag from the viscous resistance of the 3D environment. The magnitude of cell traction is proportional to the number of adhesions, *n*_*bonds*_, as:

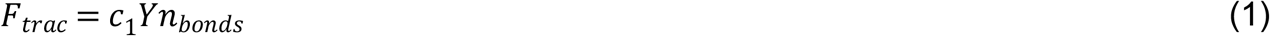

where *c*_1_ = 1 is a constant of proportionality (units of area) (31).

Environment drag, *F*_drag_, from the ECM viscous environment is:

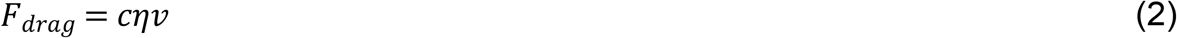

where c = 6*πr*_*cell*_ is a cell shape parameter, assuming a spherical cell of radius, *r*_*cell*_, in an infinitely viscous medium; *η* = 55 Pa-s is the effective viscosity of the medium; *v* is cell velocity and implies a velocity-dependent opposing force, associated with the viscoelastic character of the surrounding ECM (31).

Assuming force balance between *F*_*trac*_ and *F*_*drag*_, the total distance a cell can move at each timestep is computed by multiplying the velocity, *v*, by the time step. The model then evaluates a probability for migration, based on the ratio between the distance between neighboring pores in the matrix and the distance the cell can move, as: 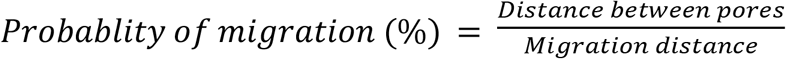. Based on the migration probability, the model evaluates each cell movement. If a cell successfully migrates to a neighboring pore, the origin pore and destination pore are updated to account for crowding during the simulation.

Following migration, cells have a chance to proliferate. In the model, cell proliferation follows a constant pre-set probability or replication rate and, in most cases, preferentially occurs following migration. However, in cases in which crowding prevents migration, cells may proliferate without migrating. Newly created daughter cells can also migrate, following the identical migration steps as the parent cell. However, unlike the parent cell, daughter cells cannot proliferate after migration.

### Adhesions clutches between cells and ECM

Integrin adhesions between each cell and the surrounding matrix surface are treated as implicit molecular clutches that dynamically bind and unbind matrix ligands, depending on the contact area between each cell and its pore. Each bond between integrin and matrix ligands has a defined stiffness, which depends upon matrix rigidity, as: 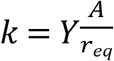, where *Y* is Young’s modulus of the ECM, *A* is the cross-sectional area of individual integrin-ligand bonds, assuming each bond has a circular shape of 10 nm radius, of the order of the radius of integrin conformation in the active state, *r*_*eq*_ = 25 nm is the equilibrium separation between cell and pore surface, corresponding to integrin headpiece extension (32). The forces on each integrin-ligand bond, *F*, is computed assuming a harmonic potential interaction, *F* = *k*(*r* − *r*_*eq*_), where *r*_*eq*_ varies according to a gaussian distribution with an average of 0.04 nm and a standard deviation of 0.001 nm to reflect different integrin activation states varying for headpiece extension. The headpiece swings from bent to extended and results in varying distances with the matrix. The lifetime of individual integrin-ligand bonds follows catch-bond kinetics (33) and depends upon the force on each integrin ligand bond, *F*, as: 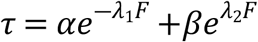.

The effective number of integrin-ligand bonds, *n*_*bonds*_, is calculated considering that the maximum number of these bonds, *n*_*max*_, is formed when all integrins are ligated and remain bound for their maximum bond lifetime, *τ*_*max*_, as *n*_*bonds*_ = *L* × *n*_*max*_*τ*/*τ*_*max*_ where *L* represents the normalized ligand concentration which ranges from 0-1 and modulates the number of effective integrin-ligand bonds; 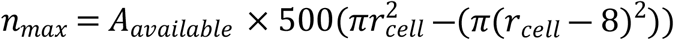, where *A*_*available*_ represents the ratio of the available pore surface area to that available in the pore itself, which accounts for the current number of cells in the pore; 500 integrins/μm^2^ are assumed to bind ligands from a portion of membrane surface mimicking the cell front and that scales with the cell radius, *r*_*cell*_.

## Results

### ECM microarchitecture and stiffness influence cell migration speed

In order to understand how the external environment affects cell migration, we first measured the migration velocity of lung adenocarcinoma cells in 3D collagen gels with different porosity and Young modulus, *Y* (Fig. 2A). In Collagen Type I 3D matrices, gelation kinetics determine matrix stiffness, fiber stiffness, and pore size. Higher gelation temperature resulted in more rapid gelation, reducing pore size and increasing material stiffness (3, 22). Collagen gels prepared at 20ºC exhibit pore size ∼10 μm, high porosity, ∼75%, and high *Y*, ∼1.5kPa, while gels prepared at 20ºC exhibit pore size ∼5 μm, low porosity, ∼65%, and low Young modulus, *Y* ∼ 1 kPa (3, 22). We found that cells in gels prepared at 20ºC migrated at 0.1290 μm/hr, 8% faster than gels prepared at 37ºC, which migrated at 0.1333 μm/hr (Fig. 2B). The increase in cell velocity on stiffer and more porous ECM is consistent with previous data from fibroblasts migration on similar polymerized rat tail collagen gels (3, 22). These results support the notion that the properties of the external environment regulate 3D cell migration speed. However, while our experiments confirm that environments with different mechanical properties and microstructure regulate 3D cell migration, they do not allow for direct evaluation of the relative contributions of *Y* and porosity on 3D cell motion.

**Figure 2.**
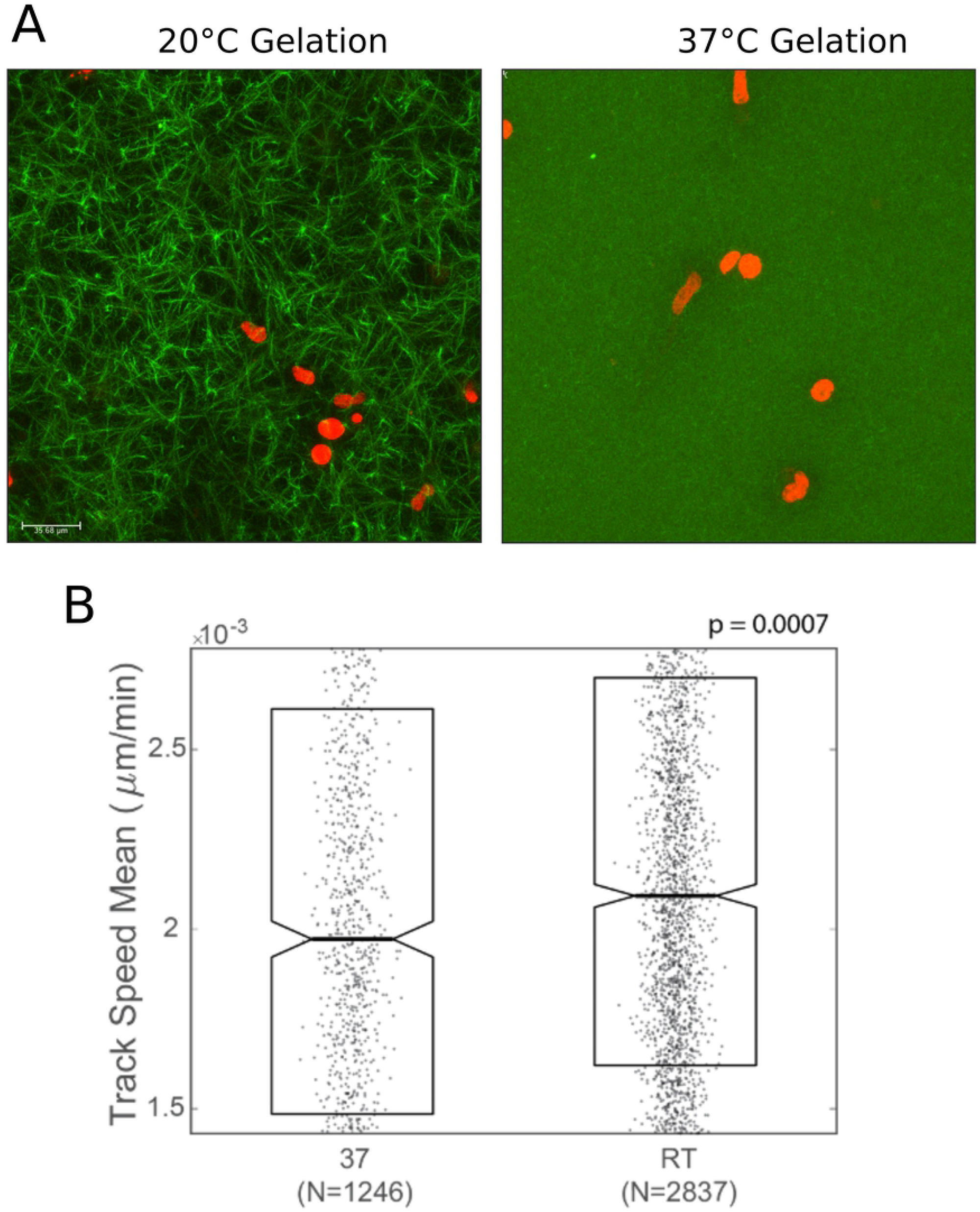
Cell invasion in collagen gels. **A**. H1299 lung adenocarcinoma cells were labeled with H_2_B-mCherry and embedded in 3-dimensional rat-tail type I collagen matrices gelled at defined temperatures. Before imaging by confocal microscopy, collagen was labeled with CNA35-GFP. Maximum intensity projections of X um sections are displayed. **B**. Boxplot of invasion speed, calculated from 14 hours of tracking the cell nuclei and calculating the average speed per track. Boxes show 50^th^ percentile and notches are 95% CI.

**Figure 3.**
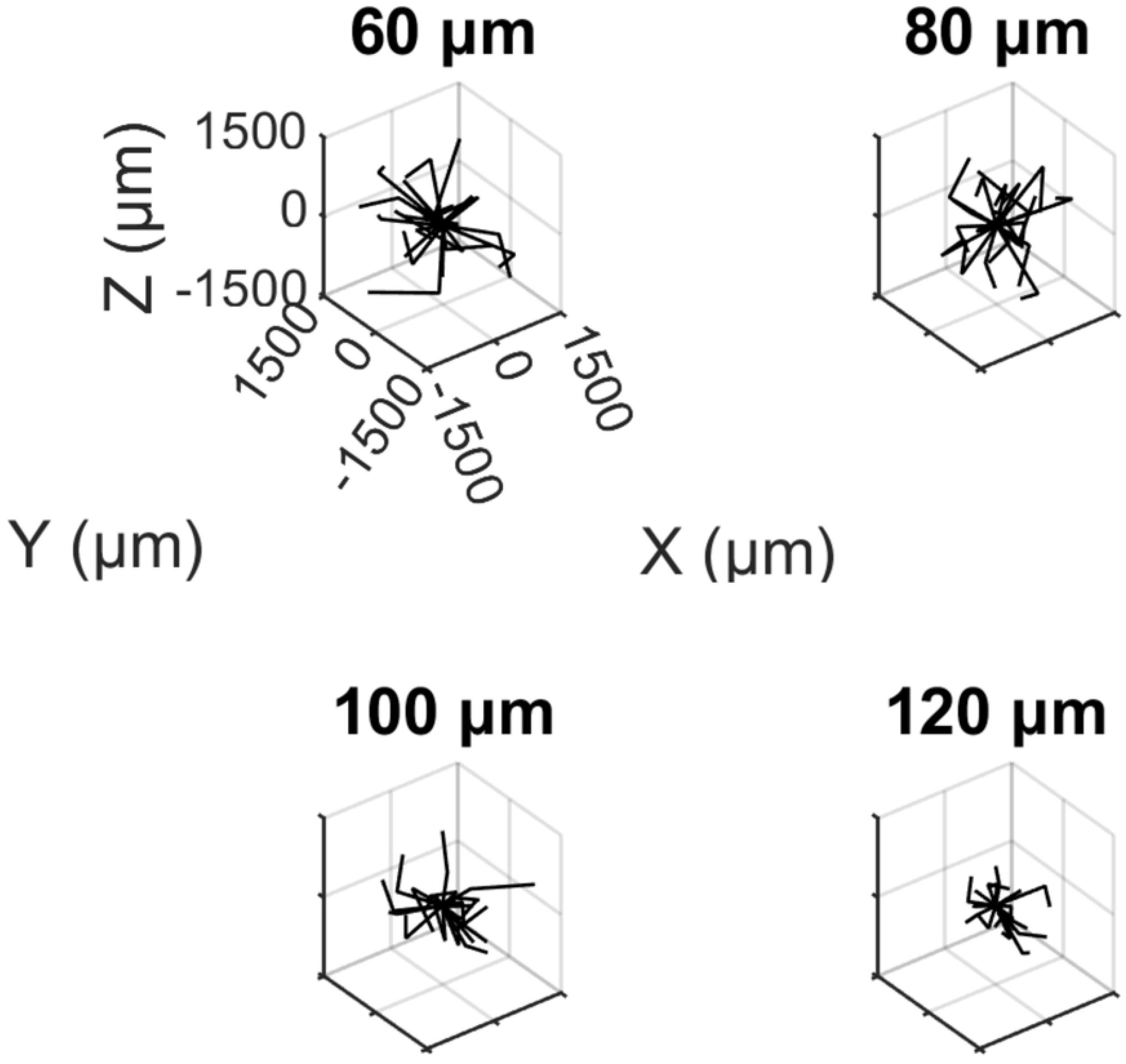
Large pores reduce 3D cell migration. Representation of the cell path using different ECM pore diameters. Simulations were run using cell density of 1.13 × 10^−6^ cells/μm^3^, *Y* = 2-4 kPa, and porosity 91%.

### Effect of environment microstructure on 3D cell migration

In order to evaluate the effect of matrix microstructure on 3D cell migration, we first ran simulations in which matrix rigidity was fixed and pore size was systematically varied. We tested pore sizes from 60 μm to 120 μm, which models the pore sizes in soft tissue and tissue scaffolds engineered for tissue regeneration and doesn’t limit migration by nuclear constriction (34). Using cell density of 1.13 × 10^−6^ cells/μm^3^, *Y* = 2-4 kPa, and porosity 91%, increasing pore size reduced the distance traveled by cells within 2000 simulated hours (Fig. 4) and decreased cell migration speed from a maximum of ∼ 200 μm/hr to a maximum of ∼ 20 μm/hr (Fig. 4A).

**Figure 4.**
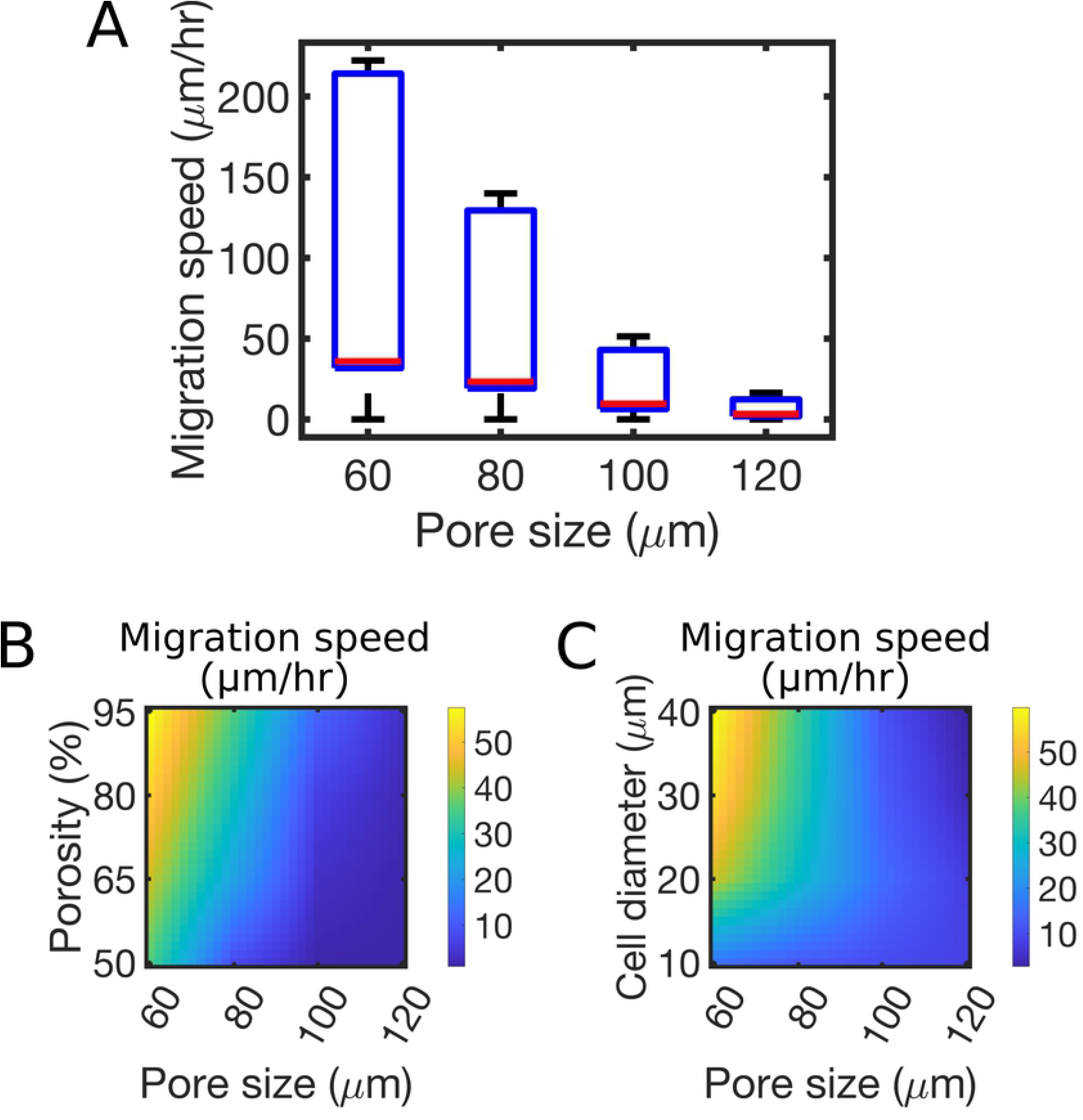
3D cell migration speed decreases with increasing ECM pore size. **A**. Boxplot of cell migration speed at varying pore sizes, from simulation runs at *Y* = 2-4 kPa, porosity 91%, and cell diameter 35 μm. Data was extracted at 2000 hours of simulations. **B**. Heatmap of average cell migration speed varying matrix pore size and porosity. Data was computed from simulation runs at *Y* = 6.3 kPa and using a fixed cell diameter of 35 μm, at 1600 hr. **C**. Heatmap of average cell migration speed varying matrix pore size and cell diameter. Data was computed from simulation runs at *Y* = 6.3 kPa, with fixed porosity of 91%, at 1600 hours of simulation.

We next tested the combined effects of pore size and porosity. Cells moved the fastest at small pore sizes and high porosities, equating to a greater number of small pores (Fig. 4B). With *Y* = 6.3 kPa and high porosity (80-95%), increasing pore size caused a rapid decrease in cell migration speed from ∼ 60 μm/hr to 10 μm/hr (Fig. 4B). With *Y* = 6.3 kPa and small pore size (60 μm), decreasing porosity from 95% to 50% decreased migration speed from ∼ 60 μm/hr to ∼ 30 μm/hr (Fig. 4B). This suggests that cell crowding limits migration. At low porosity, the same number of cells concentrate in fewer pores, so movement into neighboring pores is limited since more of the neighboring pores are full. With larger pore sizes (100 μm - 120 μm), migration occurred at a slow rate of ∼10 μm/hr and was not further reduced by decreased porosity (Fig. 4B). Thus, when pores are large and cells can make few contacts with their environment, reducing the space for this limited contact does not reduce the migration speed.

We tested the conclusion that a larger ratio between surface area available for cell-matrix connections and volume of smaller pores promotes migration by increasing the size of the modeled cells. For pore sizes below 100 μm, increasing cell diameter also enhanced migration speed using small pore size (60 μm) (Fig. 4C). Collectively, these results support the view that an increase in matrix porosity mitigates the extent to which overcrowding resulting from large cells or small pores decreases cell migration rates. Overcrowding has been reported as a factor crucial to effective cell proliferation in 3D in several previous studies (6, 35, 36).

### The model shows that 3D cell migration speed varies biphasically with environment stiffness

In order to evaluate the effect of environment stiffness on 3D cell migration, we systematically varied Young modulus, *Y*, of the lattice from 2 to 8 kPa, a range typical of soft tissues including spleen, pancreas, skin, breast, and dental pulp (1, 37, 38), as well as scaffolds for tissue engineering constructs (39). Using 91% porosity, pore diameter 60 μm and cell diameter 35 μm, cell migration speed first increased from 20 to 150 μm/hr at *Y* = 4 kPa, then decreased to almost zero at *Y* = 8 kPa (Fig. 5A). This biphasic relation between 3D cell migration speed and environment stiffness emerges from the biphasic relation between the number of adhesions and ECM stiffness (Fig. S1A), which is also reflected in the average traction force per cell (Fig. S1B).

**Figure 5.**
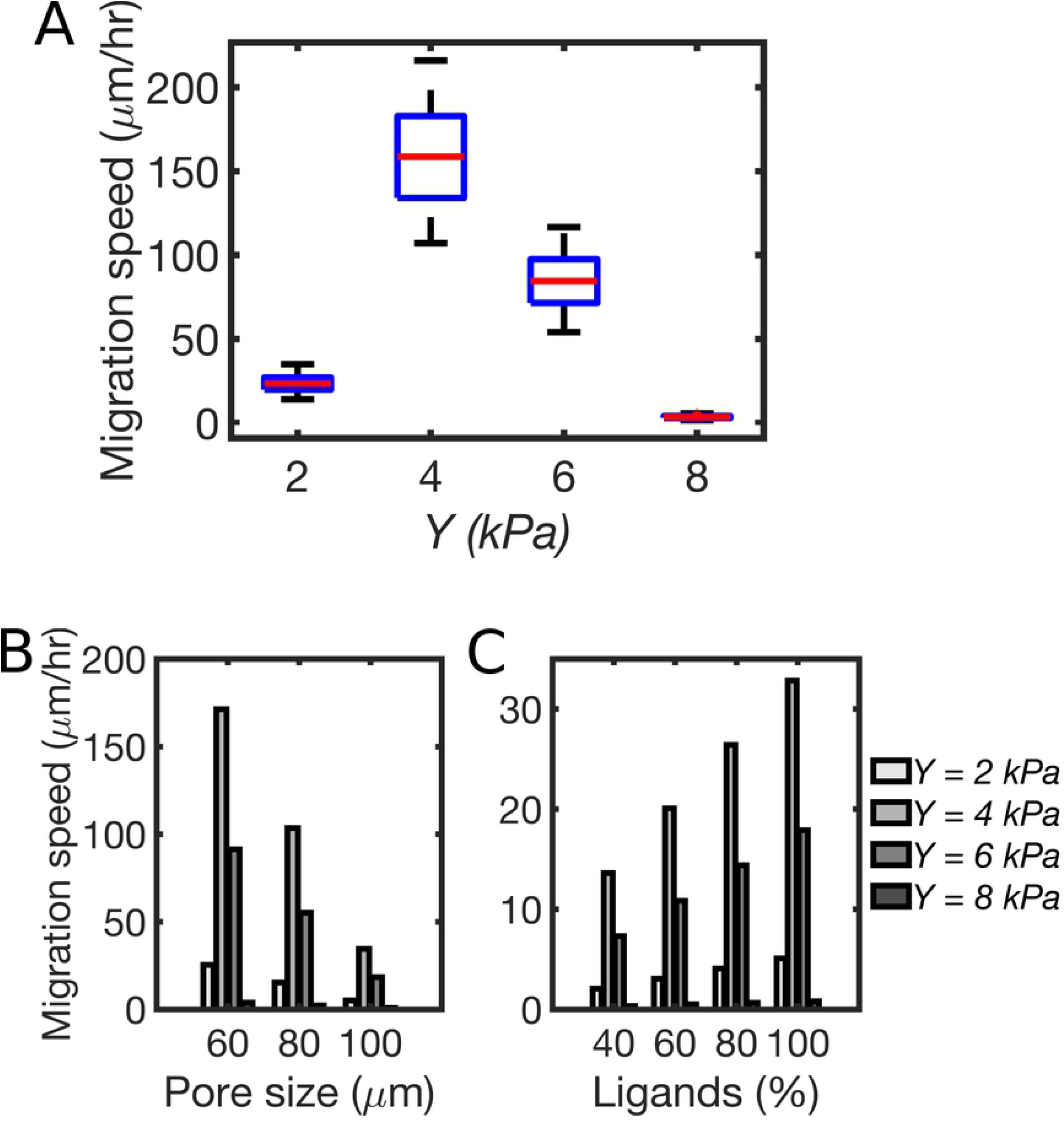
Migration speed varies biphasically with environment stiffness. **A**. Boxplot of cell migration speed for different values of matrix *Y*. Results are collected from simulation runs using porosity 91%, pore diameter 60 μm, cell diameter 35 μm, and 100% ligands. All data was calculated as averages from all cells, between 1500-2000 hr of simulations. **B**. Bar-plot of average migration speed varying pore size and matrix *Y*. Data was obtained from simulation runs using porosity 91%, cell diameter 35 μm, and 100% ligands. All data are calculated as averages from all cells at 1600 hours of simulation. **C**. Bar-plot of average migration speed varying ligand density (percentage relative to active integrins) and matrix *Y*. Data was obtained from simulation runs using porosity 91%, pore size 100 μm, cell diameter 35 μm, and cell proliferation rate 1e-7 s^-1^. All data are calculated as averages from all cells at 1600 hours of simulation.

We next sought to understand how different pore sizes affect migration speed as a function of lattice stiffness. In order to examine whether the biphasic trend of migration speed relative to stiffness was independent of pore size, we varied pore size from 60 to 100 μm, a range that presented differences in cell motion at all porosities (Fig. 4B). Migration speed decreased with increasing pore size, consistent with results in Fig. 4. While the biphasic relation between migration speed and matrix stiffness was maintained at all tested pore sizes, the magnitude of change with the different stiffnesses decreased with increasing pore size (Fig. 5B). This finding is consistent with changes in adhesion number and traction force with pore size, which also present a biphasic relation with stiffness (Fig. S1A-B).

Since pore size reduced migration speed and reduced the effect of the biphasic relationship between matrix stiffness and migration speed, we hypothesized that ligand density controls migration speed via the molecular clutch mechanism. Ligand density determines the maximum number of adhesions that a cell can form with the lattice (see Methods). The molecular clutch mechanism determines the number of adhesions each cell forms with the lattice pore, based on the force acting on them (see Methods). We tested if ligand density controls cell migration speed by reducing the probability of integrin ligation as a proxy for reduced ligand density. A reduction of ligand density from 100% (meaning that all integrins become ligated) to 40% (less than half integrins become ligated) corresponds to a proportional decrease in cell migration speed (Fig. 5C). Using cells with larger diameters shifts the migration speed towards higher values at all ligand densities, because of the larger surface area between cell and pore (Fig. S1C). Accordingly, systematic variations of cell diameter or ligand density show a biphasic relation of cell speed versus *Y*, where the peak is proportional to the relative cell surface (Fig. S2A) and the number of adhesions, respectively (Fig. S2B). Collectively, these results support the existence of a biphasic relation between cell migration speed and matrix stiffness in different conditions of environment microstructure and ligand density. The relation between cell migration speed and matrix stiffness has magnitude proportional to the number of adhesions and cell traction force. The peak migration speed corresponds to the stiffness at which the maximum number of adhesions is allowed, which is also modulated by pore size (Fig. S1). This further indicates that the amount of surface area available for the cells to bind, together with ligand density, determines the effective number of possible adhesions mediating cell motion in 3D.

### How cell proliferation affects 3D cell migration in different environments

We next tested the effect of cell proliferation on 3D cell migration. Using a physiological proliferation rate of 10^−7^ s^-1^ and increasing pore size from 60 to 120 μm, our model shows a reduction in average cell speed from 65 μm/hr to 10 μm/hr (Fig. 6A). This results from the larger ratio between surface area and volume of smaller pores, allowing cells to form more adhesions for the same number of cells in the pore. Additionally, this is consistent with previous experiments on mouse fibroblasts in collagen-glycosaminoglycan scaffolds (5). More generally, using pore sizes smaller than 100 μm, systematic variations in proliferation rate from 1 to 2.5 10^−7^ s^-1^ reduced average migration speed from ∼ 65 μm/hr to ∼ 55 μm/hr (Fig. 6A). At all proliferation rates, using variations in stiffness from 2 to 8 kPa, a peak in migration speed was consistently observed for 4 kPa, (Fig. 6B), similar to runs with fixed cell density (no proliferations) (Fig. 5A). However, both the maximum migration speed and the spread of this peak as a function of the proliferation rate decreased with increasing proliferation rate (Fig. 6B and Fig. S3A-B). Smaller pores enhanced cell migration at all proliferation rates (Fig. S3C). Increasing the proliferation rate increases the overall cell density, thereby preventing cells from migrating to neighboring pores due to crowding. As a result, we expect the overall migration rate to decrease.

**Figure 6.**
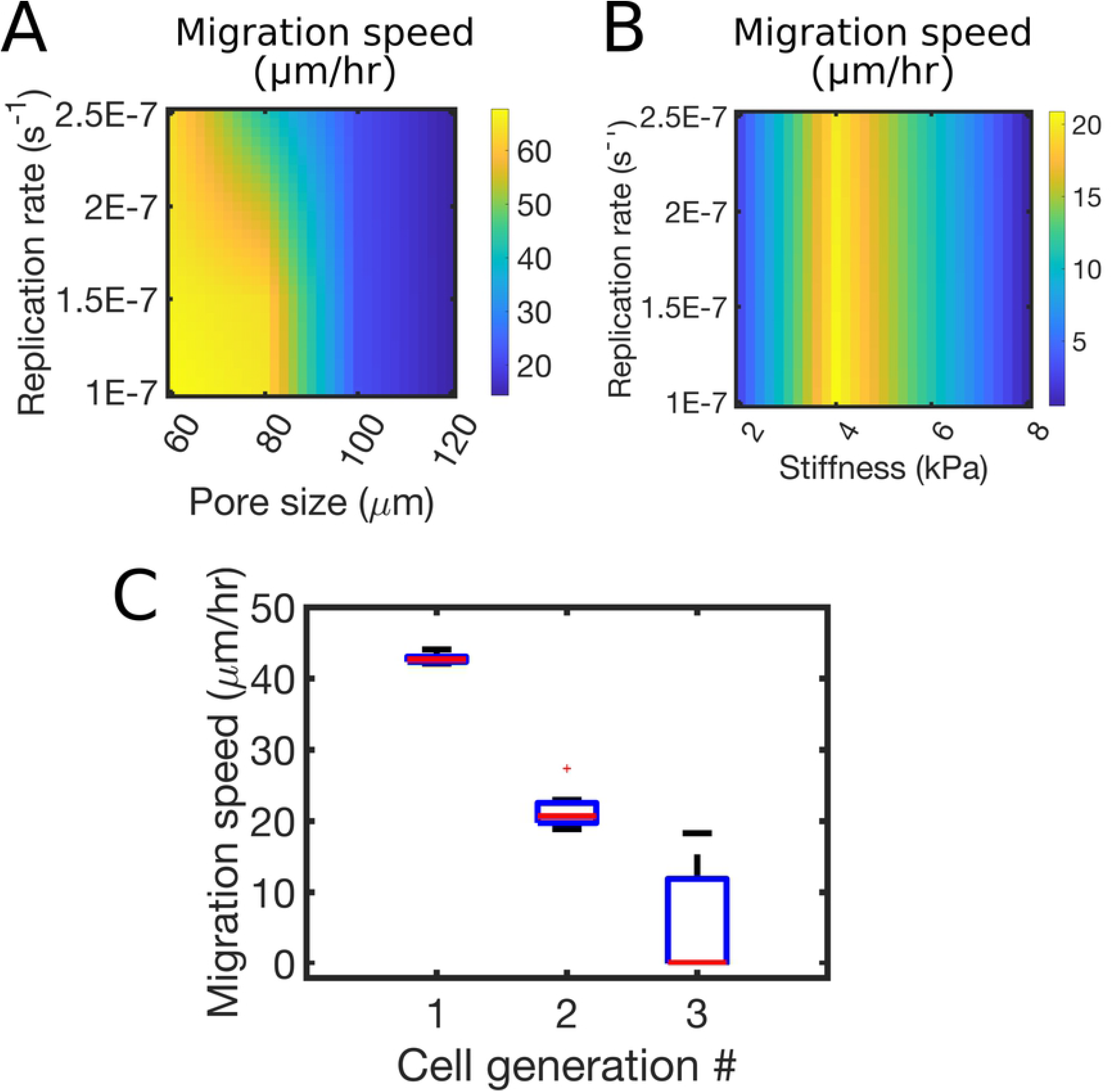
Cell proliferation reduces 3D migration speed. **A**. Heatmap of average migration speed varying cell proliferation rate and matrix pore size. Data are extracted from all cells in the system, at 1600 hours, from simulation runs using *Y* = 6.3 kPa, porosity 91%, and cell diameter 35 μm. **B**. Heatmap of average migration speed varying cell proliferation rate and matrix *Y*. Data are extracted from all cells in the system, at 1900 hr, from simulation runs using pore size 100 μm, porosity 91%, and cell diameter 35 μm. **C**. Boxplot of cell migration speed per cell generation, using *Y* = 4 kPa, porosity 91%, pore size 100 μm, and cell diameter 35 μm. Results are collected from all cells of the corresponding generation, between 300 and 1500 hours of simulations.

In order to better understand why cell proliferation decreased cell migration speed, we analyzed the distribution of velocities for different cell generations. The distribution of cell migration speed shifted towards lower values by increasing cell generation (Fig. 6C), indicating that crowding effects limit cell migration. Quantification of the fraction of immotile cells at different proliferation rates showed that cells that cease moving, decrease with increasing stiffness, up to 4 kPa, then increase (Fig. S3E). This corresponds to the first increase in cell speed, followed by decreases as a function of stiffness (Fig. S3D). When cells proliferate, the matrix becomes crowded, and cells do not find space to move to the neighboring lattice points. This inhibited their displacement, which decreased overall migration speed. Collectively, these data indicate that increased cell proliferation decreases migration but does not affect how this speed varies neither as a function of the matrix microstructure nor its mechanical properties. In fact, the decrease of cell speed with increasing pore size and its biphasic relation with matrix stiffness is maintained at all tested proliferation rates (Fig. 6A-B).

## Discussion

The mechanisms underlying 3D cell motion remain elusive. In this study, we developed a computational model of 3D cell migration to understand how the ECM regulates migration speed through adhesion clutches. The model is highly flexible and is able to characterize the migratory behavior of cells in several conditions of matrix rigidity and microstructure, as well as using different values of cell proliferation. In the simulations, characteristics like cell directionality and adhesion number are not imposed a priori; they are the result of the interaction with the matrix and force-feedbacks between the actomyosin cytoskeleton of the cells and their external environment. As an outcome, we focus on experimentally addressable characteristics of cell locomotion, i.e., cell speed, predicting how this is influenced by manipulations of either cell or matrix properties.

Collectively, the results from the model indicate that mechanical factors are not responsible for the observed effect of scaffold pore size and porosity on cell motility. At all tested stiffnesses, cell migration speed decreases with increasing pore size and increases with porosity (Fig. 3). Increasing pore size decreases the matrix surface area to volume ratio, which reduces the number of adhesions that cells can create, as observed in previous experiments on fibroblasts in collagen hydrogels (7). The positive correlation between porosity and cell migration speed is consistent with experiments on human dermal fibroblasts in 3D silk fibroin scaffolds and studies on breast cancer cells in tunable gelatin matrices (6, 40). Additionally, other experiments, including studies of fibroblasts motion in scaffolds, have previously found a positive relationship between smaller pore sizes and migratory speeds (5, 19, 41, 42), which align with the results from our simulations. Collectively, these similar trends regarding 3D cell migration relative to variations in porosity and pore size point to the importance of porosity and pore size in modulating crowding and in regulating the dynamics of adhesion clutches for 3D cell motion.

Our model implements a lattice point approach with clutch-dependent cell proliferation and migration. The lattice point approach was used in previous models to simulate cell proliferation and migration, but differently and with an another objective. The objective of previous lattice point models of 3D cell migration was to find the best design parameters for 3D scaffolds for tissue regeneration (26, 27). Hence the focus was on mechanosensitive cell proliferation, where migration was an emergent property. By contrast, the goal of our model is to understand fundamentally how cells move in 3D. Hence the algorithm treats migration first and differentiation is an emergent property. Our model uses differentiation as a constant rate, but proliferations is also contingent on successful catch-bond-based migration and local crowding conditions; thus it emerges indirectly as a mechanosensory behavior from migration.

Regulation of cell migration through catch-bond mechanics through the motor-clutch model has only previously been simulated in 2D (43), but recent experiments have highlighted the existence of adhesion clutches in 3D (22, 29, 30). For the first time, our model incorporates the dynamics of mechanosensitive clutch-based adhesions in 3D to study their effects on cell migratory behavior in ECM different for microstructure and mechanics. The molecular clutch-based dynamics are responsible for the effect of 3D matrix stiffness on cell migratory rates, which exhibit an increase then a decrease (biphasic trend) with increasing stiffness, consistent with results seen in 2D (23–25). The adhesion clutch dynamics can thus underlie the biphasic relation between 3D cell invasion speed and environment mechanics that has been observed in several experimental systems, including tumor cells migration in polyacrylamide-based hydrogels (19), myoblasts in PLLA/PGLA scaffolds (39), and mouse fibroblasts in collagen-glycosaminoglycan scaffolds (5).

Our model doesn’t incorporate information regarding the change in the shape of the cell or constrain on nuclear passage as the cell moves through the ECM. It has been reported that due to the steric resistance of the matrix, a cell might change its shape to migrate through the ECM and that small pores can limit nuclear passage (3, 44, 45). Also, a detailed description of the chemical basis of actomyosin contraction is not addressed in our model. Such aspects of cell migration are beyond the scope of our model at this moment. In the future, features that may be incorporated in our model include cell compressibility, nuclear dimension and also the notion of dual pore sizes within the same ECM. Dual pore sizes or porosities center around the notion that smaller pores facilitate nutrient supply, waste removal, and cell proliferation, while larger pores encourage proliferation and interconnectivity (46).

To conclude, our model supports a picture in which cells respond to the effective microstructure of the environment through adhesion clutches working as local matrix attachment points that regulate cell motion through molecular force-dependent activities. Notably, the matrix microstructure impacts 3D cell motion independently from mechanical factors. These results highlight the important role the molecular clutch may have on cell migration in 3D. By better understanding how stiffness and microstructure affect cell motility through mechanosensitive adhesion clutches, we can design better treatments, scaffolds for tissue regeneration, and therapies that provide ideal environments that may enhance or mitigate cell migration and replication.

## Acknowledgements

TCB was supported by NSF BMMB 2044394. MCM was supported by American Cancer Society RSG CSM130435.

## Supplementary information

### Model initialization

Before generating the 3D domain, seeding the cells in the domain, and running the simulations, the model requires parameters corresponding to three categories: 3D environment, cell properties, and simulation parameters. The input parameters for the 3D environment include side dimension of the simulated cube (in *μm*), porosity (*%*), pore size or diameter (in *μm*), cell packing density (%), and Young’s modulus of the material, *Y*. The input cell parameters include cell diameter (*μm*), initial cell density, and the rate of cell replication (*s*^*-1*^). Simulation parameters include time step, total simulation time, and frequency of data recording.

### Generation of the 3D Environment

The 3D environment is generated as an object from a class known as *Scaffold*. This class stores and keeps track of the properties of the 3D environment. To generate the lattice, the total 3D volume, including empty and non-empty spaces, *V*_*matrix*_, is calculated by raising the input side dimension of the environment to the third power:

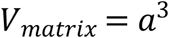

The total volume that is “porous” (empty) is calculated by multiplying the input percentage of porosity by *V*_*matrix*_. Therefore, given a specific pore size, the porosity determines the total empty (porous) volume in the lattice, as:

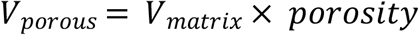

The volume of each pore, *V*_*pore*_, is calculated from the pore diameter, *d*_*pore*_, with the assumption that the pore is spherical:

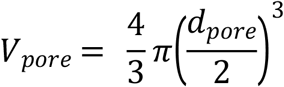

The number of pores in the 3D matrix is then calculated by dividing the total empty volume *V*_*porous*_, by the volume of each pore, *V*_*pore*_, and taking the lower bound value:

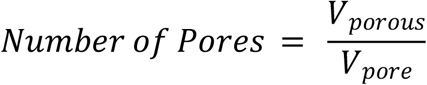

The number of pores per side length is then calculated as the lower bound cube root of the number of pores:

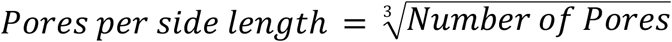

In the code, each pore is indexed from a list using three indices. Each index corresponds to X, Y, and Z coordinates depending on the position of the pore in the lattice.

The maximum occupancy of cells in each pore is calculated from the volume of each cell using the cell diameter, *d*_*cell*_:

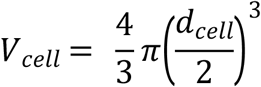

The maximum number of cells in a pore is calculated by dividing *V*_*pore*_ by *V*_*cell*_ multiplied the *packing density* and rounded down:

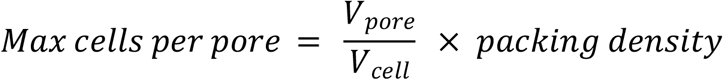

The 3D matrix is initially seeded with cells randomly throughout all the pores in the matrix.

### Data Generation

Based on the writing frequency specified in the program, seven different characteristics of each cell are recorded and stored in a simulation comma-separated values (CSV) file. These eight characteristics are time (hours), unique cell ID, cell generation, number of cell moves, instantaneous traction force, X-coordinates, Y-coordinates, and Z-coordinates. All data-derived figures were generated from the use of these data.

## Supplementary Figures

**Figure S1. Cell Adhesions and traction force vary biphasically with matrix stiffness. A**. The average number of adhesions as a function of pore size for different matrix rigidities. **B**. Average traction force per cell varying pore size and matrix rigidity. C. Average migration velocity for cells with different diameters (35 and 50 μm) varying the percentage of ligands. Data are collected from runs using 3D matrices with a cell density of 1.13 × 10^−6^ cells/μm^3^ with no proliferation and 91% porosity and at 1000 hours of simulation.

**Figure S2. Cell speed depends on cell size and ligand density. A**. Heatmap of average cell speed varying cell diameter and *Y*. **B**. Average number of adhesions per cell varying *Y* and ligand density. Data are collected at 2000 hr of simulations using 91% porosity.

**Figure S3. Cell proliferation regulates migration speed through crowding. A**. Heatmap of average cell migration speed varying lattice pore size and *Y*, without cell proliferation. **B**. Heatmap of average cell migration speed varying lattice pore size and *Y*, with cell proliferation rate 10^−7^ s^-1^. **C**. Heatmap of average cell migration speed by varying cell proliferation rate and lattice pore size, using *Y* = 6.3 kPa and porosity 91%. D. Average cell migration speed varying cell proliferation rate and *Y*, using pore fixed size 100 μm. **E**. Corresponding immobile fraction of cells relative to the total number of cells. Data are collected from 2000 hr of simulations.

## Notes

### Competing Interest Statement

The authors have declared no competing interest.

## References

1. van Helvert, S., C. Storm, and P. Friedl. 2018. Mechanoreciprocity in cell migration. Nat. Cell Biol. 20: 8–20.

2. Riching, K.M., B.L. Cox, M.R. Salick, C. Pehlke, A.S. Riching, S.M. Ponik, B.R. Bass, W.C. Crone, Y. Jiang, A.M. Weaver, K.W. Eliceiri, and P.J. Keely. 2014. 3D collagen alignment limits protrusions to enhance breast cancer cell persistence. Biophys. J. 107: 2546–2558.

3. Wolf, K., M. te Lindert, M. Krause, S. Alexander, J. te Riet, A.L. Willis, R.M. Hoffman, C.G. Figdor, S.J. Weiss, and P. Friedl. 2013. Physical limits of cell migration: Control by ECM space and nuclear deformation and tuning by proteolysis and traction force. J. Cell Biol. 201: 1069–1084.

4. Tien, J., U. Ghani, Y.W. Dance, A.J. Seibel, M.Ç. Karakan, K.L. Ekinci, and C.M. Nelson. 2020. Matrix Pore Size Governs Escape of Human Breast Cancer Cells from a Microtumor to an Empty Cavity. iScience. 23: 101673.

5. Harley, B.A.C., H. Do Kim, M.H. Zaman, I. V. Yannas, D.A. Lauffenburger, and L.J. Gibson. 2008. Microarchitecture of three-dimensional scaffolds influences cell migration behavior via junction interactions. Biophys. J. 95: 4013–4024.

6. Mandal, B.B., and S.C. Kundu. 2009. Cell proliferation and migration in silk fibroin 3D scaffolds. Biomaterials. 30: 2956–2965.

7. Gun’ko, V.M., L.I. Mikhalovska, I.N. Savina, R. V Shevchenko, S.L. James, P.E. Tomlins, and S. V Mikhalovsky. 2010. Characterisation and performance of hydrogel tissue scaffolds. Soft Matter. 6: 5351–5358.

8. Mierke, C.T., B. Frey, M. Fellner, M. Herrmann, and B. Fabry. 2011. Integrin α5β1 facilitates cancer cell invasion through enhanced contractile forces. J. Cell Sci. 124: 369–383.

9. Chiu, C.-L., J.S. Aguilar, C.Y. Tsai, G. Wu, E. Gratton, and M.A. Digman. 2014. Nanoimaging of Focal Adhesion Dynamics in 3D. PLoS One. 9: e99896.

10. Plotnikov, S. V., A.M. Pasapera, B. Sabass, and C.M. Waterman. 2012. Force fluctuations within focal adhesions mediate ECM-rigidity sensing to guide directed cell migration. Cell.

11. Doyle, A.D., D.J. Sykora, G.G. Pacheco, M.L. Kutys, and K.M. Yamada. 2021. 3D mesenchymal cell migration is driven by anterior cellular contraction that generates an extracellular matrix prestrain. Dev. Cell. 56: 826-841.e4.

12. Gupton, S.L., and C.M. Waterman-Storer. 2006. Spatiotemporal feedback between actomyosin and focal-adhesion systems optimizes rapid cell migration. Cell. 125: 1361–1374.

13. Hu, K., L. Ji, K.T. Applegate, G. Danuser, and C.M. Waterman-Storer. 2007. Differential transmission of actin motion within focal adhesions. Science. 315: 111–115.

14. Lele, T.P., C.K. Thodeti, and D.E. Ingber. 2006. Force meets chemistry: Analysis of mechanochemical conversion in focal adhesions using fluorescence recovery after photobleaching. J. Cell. Biochem. 97: 1175–1183.

15. DuChez, B.J., A.D. Doyle, E.K. Dimitriadis, and K.M. Yamada. 2019. Durotaxis by Human Cancer Cells. Biophys. J. 116: 670–683.

16. Lo, C.M., H.B. Wang, M. Dembo, and Y.L. Wang. 2000. Cell movement is guided by the rigidity of the substrate. Biophys. J. 79: 144–52.

17. Raab, M., J. Swift, P.C.D.P. Dingal, P. Shah, J.W. Shin, and D.E. Discher. 2012. Crawling from soft to stiff matrix polarizes the cytoskeleton and phosphoregulates myosin-II heavy chain. J. Cell Biol.

18. Elosegui-Artola, A., R. Oria, Y. Chen, A. Kosmalska, C. Pérez-González, N. Castro, C. Zhu, X. Trepat, and P. Roca-Cusachs. 2016. Mechanical regulation of a molecular clutch defines force transmission and transduction in response to matrix rigidity. Nat. Cell Biol.

19. Pathak, A., and S. Kumar. 2012. Independent regulation of tumor cell migration by matrix stiffness and confinement. Proc. Natl. Acad. Sci. U. S. A. 109: 10334–10339.

20. Lang, N.R., K. Skodzek, S. Hurst, A. Mainka, J. Steinwachs, J. Schneider, K.E. Aifantis, and B. Fabry. 2015. Biphasic response of cell invasion to matrix stiffness in three-dimensional biopolymer networks. Acta Biomater. 13: 61–67.

21. Zaman, M.H., L.M. Trapani, A.L. Sieminski, A. Siemeski, D. Mackellar, H. Gong, R.D. Kamm, A. Wells, D.A. Lauffenburger, and P. Matsudaira. 2006. Migration of tumor cells in 3D matrices is governed by matrix stiffness along with cell-matrix adhesion and proteolysis. Proc. Natl. Acad. Sci. U. S. A. 103: 10889–94.

22. Doyle, A.D., N. Carvajal, A. Jin, K. Matsumoto, and K.M. Yamada. 2015. Local 3D matrix microenvironment regulates cell migration through spatiotemporal dynamics of contractility-dependent adhesions. Nat. Commun. 6: 8720.

23. Bangasser, B.L., G.A. Shamsan, C.E. Chan, K.N. Opoku, E. Tüzel, B.W. Schlichtmann, J.A. Kasim, B.J. Fuller, B.R. McCullough, S.S. Rosenfeld, and D.J. Odde. 2017. Shifting the optimal stiffness for cell migration. Nat. Commun. 8: 1–10.

24. Bangasser, B.L., and D.J. Odde. 2013. Master equation-based analysis of a motor-clutch model for cell traction force. Cell. Mol. Bioeng. 6: 449–459.

25. Oakes, P.W., T.C. Bidone, Y. Beckham, A.V. Skeeters, G.R. Ramirez-San Juan, S.P. Winter, G.A. Voth, and M.L. Gardel. 2018. Lamellipodium is a myosin-independent mechanosensor. Proc. Natl. Acad. Sci. U. S. A. 115.

26. Byrne, D.P., D. Lacroix, J.A. Planell, D.J. Kelly, and P.J. Prendergast. 2007. Simulation of tissue differentiation in a scaffold as a function of porosity, Young’s modulus and dissolution rate: application of mechanobiological models in tissue engineering. Biomaterials. 28: 5544–5554.

27. Checa, S., and P.J. Prendergast. 2009. A mechanobiological model for tissue differentiation that includes angiogenesis: a lattice-based modeling approach. Ann. Biomed. Eng. 37: 129–145.

28. Sandino, C., S. Checa, P.J. Prendergast, and D. Lacroix. 2010. Simulation of angiogenesis and cell differentiation in a CaP scaffold subjected to compressive strains using a lattice modeling approach. Biomaterials. 31: 2446–2452.

29. Cantini, M., H. Donnelly, M.J. Dalby, and M. Salmeron-Sanchez. 2020. The Plot Thickens: The Emerging Role of Matrix Viscosity in Cell Mechanotransduction. Adv. Healthc. Mater. 9: 1901259.

30. Owen, L.M., A.S. Adhikari, M. Patel, P. Grimmer, N. Leijnse, M.C. Kim, J. Notbohm, C. Franck, and A.R. Dunn. 2017. A cytoskeletal clutch mediates cellular force transmission in a soft, three-dimensional extracellular matrix. Mol. Biol. Cell. 28: 1959–1974.

31. Zaman, M.H., R.D. Kamm, P. Matsudaira, and D.A. Lauffenburger. 2005. Computational model for cell migration in three-dimensional matrices. Biophys. J. 89: 1389–1397.

32. Xu, X.-P., E. Kim, M. Swift, J.W. Smith, N. Volkmann, and D. Hanein. 2016. Three-Dimensional Structures of Full-Length, Membrane-Embedded Human α(IIb)β(3) Integrin Complexes. Biophys. J. 110: 798–809.

33. Kong, F., A.J. García, A.P. Mould, M.J. Humphries, and C. Zhu. 2009. Demonstration of catch bonds between an integrin and its ligand. J. Cell Biol.

34. Jenkins, T.L., and D. Little. 2019. Synthetic scaffolds for musculoskeletal tissue engineering: cellular responses to fiber parameters. npj Regen. Med. 4: 15.

35. Pereira, R.C., R. Santagiuliana, L. Ceseracciu, D.P. Boso, B.A. Schrefler, and P. Decuzzi. 2020. Elucidating the Role of Matrix Porosity and Rigidity in Glioblastoma Type IV Progression. Appl. Sci.. 10.

36. Li, J., Y.T. Chong, C.P. Teng, J. Liu, and F. Wang. 2021. Microporosity mediated proliferation of preosteoblast cells on 3D printed bone scaffolds. Nano Sel. n/a.

37. Liu, J., H. Zheng, P.S.P. Poh, H.-G. Machens, and A.F. Schilling. 2015. Hydrogels for Engineering of Perfusable Vascular Networks. Int. J. Mol. Sci. 16: 15997–16016.

38. Guimarães, C.F., L. Gasperini, A.P. Marques, and R.L. Reis. 2020. The stiffness of living tissues and its implications for tissue engineering. Nat. Rev. Mater. 5: 351–370.

39. Levy-Mishali, M., J. Zoldan, and S. Levenberg. 2008. Effect of Scaffold Stiffness on Myoblast Differentiation. Tissue Eng. Part A. 15: 935–944.

40. Huang, D., Y. Nakamura, A. Ogata, and S. Kidoaki. 2020. Characterization of 3D matrix conditions for cancer cell migration with elasticity/porosity-independent tunable microfiber gels. Polym. J. 52: 333–344.

41. Karageorgiou, V., and D. Kaplan. 2005. Porosity of 3D biomaterial scaffolds and osteogenesis. Biomaterials. 26: 5474–5491.

42. Zhang, Z.-Z., D. Jiang, J.-X. Ding, S.-J. Wang, L. Zhang, J.-Y. Zhang, Y.-S. Qi, X.-S. Chen, and J.-K. Yu. 2016. Role of scaffold mean pore size in meniscus regeneration. Acta Biomater. 43: 314–326.

43. Chan, C.E., and D.J. Odde. 2008. Traction Dynamics of Filopodia on Compliant Substrates. Science (80-.). 322: 1687–1691.

44. Friedl, P., and E.-B. Bröcker. 2000. The biology of cell locomotion within three-dimensional extracellular matrix. Cell. Mol. Life Sci. C. 57: 41–64.

45. Wolf, K., I. Mazo, H. Leung, K. Engelke, U.H. von Andrian, E.I. Deryugina, A.Y. Strongin, E.-B. Bröcker, and P. Friedl. 2003. Compensation mechanism in tumor cell migration: mesenchymal-amoeboid transition after blocking of pericellular proteolysis. J. Cell Biol. 160: 267–277.

46. Rasoulianboroujeni, M., N. Kiaie, F.S. Tabatabaei, A. Yadegari, F. Fahimipour, K. Khoshroo, and L. Tayebi. 2018. Dual Porosity Protein-based Scaffolds with Enhanced Cell Infiltration and Proliferation. Sci. Rep. 8: 14889.

47. Ilina, O., P.G. Gritsenko, S. Syga, J. Lippoldt, C.A.M. La Porta, O. Chepizhko, S. Grosser, M. Vullings, G.-J. Bakker, J. Starruß, P. Bult, S. Zapperi, J.A. Käs, A. Deutsch, and P. Friedl. 2020. Cell-cell adhesion and 3D matrix confinement determine jamming transitions in breast cancer invasion. Nat. Cell Biol. 22: 1103–1115.

